# Biophysical mechanisms governing large-scale brain network dynamics underlying individual-specific variability of perception

**DOI:** 10.1101/819896

**Authors:** G. Vinodh Kumar, Shrey Dutta, Siddharth Talwar, Dipanjan Roy, Arpan Banerjee

## Abstract

Perception necessitates interaction amongst neuronal ensembles, the dynamics of which can be conceptualized as the emergent behavior of coupled dynamical systems. Here, we propose a detailed neurobiologically realistic model that captures the neural mechanisms of inter-individual variability observed in cross-modal speech perception. From raw EEG signals recorded from human participants when they were presented with speech vocalizations of McGurk-incongruent and congruent audio-visual (AV) stimuli, we computed the global coherence metric to capture the neural variability of large-scale networks. We identified that participants’ McGurk susceptibility was negatively correlated to their alpha-band global coherence. The proposed biophysical model conceptualized the global coherence dynamics emerge from coupling between the interacting neural masses - representing the sensory specific auditory/visual areas and modality non-specific associative/integrative regions. Subsequently, we could predict that an extremely weak direct AV coupling result in a decrease in alpha band global coherence - mimicking the cortical dynamics of participants with higher McGurk susceptibility. Source connectivity analysis also showed decreased connectivity between sensory specific regions in participants more susceptible to McGurk effect, thus establishing an empirical validation to the prediction. Overall, our study provides an outline to link variability in structural and functional connectivity metrics to variability of performance that can be useful for several perception & action task paradigms.

## Introduction

Neurobiologically realistic dynamical models of perception have provided attractive explanation of behavior (Moreno-Bote et al 2007, Thakur et al., 2016; Cuppini et al., 2017). Concurrently, they also provide insights to brain dynamics observed at different scales e.g., microscopic level of single neurons (Deco et al., 2007;) and macroscopic level of field recordings (David & Friston, 2003; Spiegler et al., 2011; Onojima et al., 2018). How does this framework adapt to perceptual variability across individuals during the perception of complex signals? One intuitive approach to explain perceptual variability involves characterization of stochastic fluctuations in component neural systems. Thereafter, probability distributions set by internal processes that generate prediction can follow Bayes’ optimization principles to set up perceptual object identification (Körding et al., 2007). One drawback of these descriptive Bayesian brain hypotheses is that, unlike predictive coding formulations, they are agnostic to the neurobiological mechanisms and systems-level interactions that govern perception (Ursino et al., 2017). In this article, we employ McGurk effect (McGurk & Macdonald, 1976) to demonstrate how the systems-level interactions in multisensory and unisensory subsystems can explain differences in McGurk susceptibility across individuals.

The fact that speech perception is fundamentally driven by visual cues is demonstrated in this paradigm - an auditory /*ba*/ dubbed onto visual lip movements of /*ga*/ is perceived by the listener as a completely different syllable /*da*/ (McGurk & Macdonald, 1976). Variability in the perception of the illusion has enabled researchers to unravel the neural correlates of illusory/cross-modal percept in terms of cortical activations and neural oscillations (Nath & Beauchamp, 2011; Keil et al., 2012). However, these studies have overlooked the substantial variability of McGurk susceptibility across individuals that could influence the objective behavioral and neural estimates. From a neurophysiological standpoint, how do we reconcile the neural correlates of McGurk percept under the purview of inter-individual variability or vice versa? Moreover, it raises a fundamental question on what constitutes an appropriate neural correlate. For instance, activity in superior temporal sulcus (STS) and its functional connectedness to sensory specific and fronto-parietal regions has been shown to subserve multisensory speech perception and McGurk susceptibility (Nath & Beauchamp, 2012, Keil et al., 2012) as well. Moreover, other research areas like face processing (Haxby et al., 2000), motion perception (Puce & Perrett, 2003) and social perception (Saxe, 2006) attribute activations in the STS to the particular behavior being studied. This suggests that the STS is a prime nexus for information processing; however, it may not be a neural correlate of any specific processing component.

Emerging studies indicate complex cognitive capabilities to emerge as a result of interactions amongst large-scale networks of cortical areas (Varela et al., 2001; Bressler & Menon, 2010). Therefore, a measure that could capture the ‘dynamic’ and ‘preferential’ interactions amongst distributed neuronal assemblies/brain regions will offer the most pertinent representation of the distinct neural machinery supporting McGurk susceptibility.

Recent studies have indicated that beyond a specific region of interest, a large-scale network of oscillatory brain networks underwrite the McGurk effect (Kumar et al., 2016; Kumar et al., 2017). A key question emerges, how robust is this network across individuals and whether the organization of such networks is contingent on the stimulus configurations. Secondly, what are the neural mechanisms that give rise to such network level correlates? While the first question needs to be answered empirically using a detailed neurophysiological study of underlying brain networks, the more broader question of systems-level insight require a neurobiologically inspired large-scale computational model.

We hypothesized that variability in McGurk susceptibility across individuals stem from implicit differences in their large–scale functional connectivity. In order to test that, we conducted psychophysical experiments in tandem with EEG recordings to characterize the large-scale oscillatory modes in whole-brain dynamics. Subsequently we constructed a neural mass model for multisensory perception - an abstraction of summed synapto-dendritic activity of thousands of neurons, comprising three nodes representing sensory and multisensory areas. Finally, we validated the plausible neural mechanisms made by the model using source space analysis (**Figure 1**), thus providing a link between functional metrics and underlying neurophysiological complexity.

**Figure 1:**
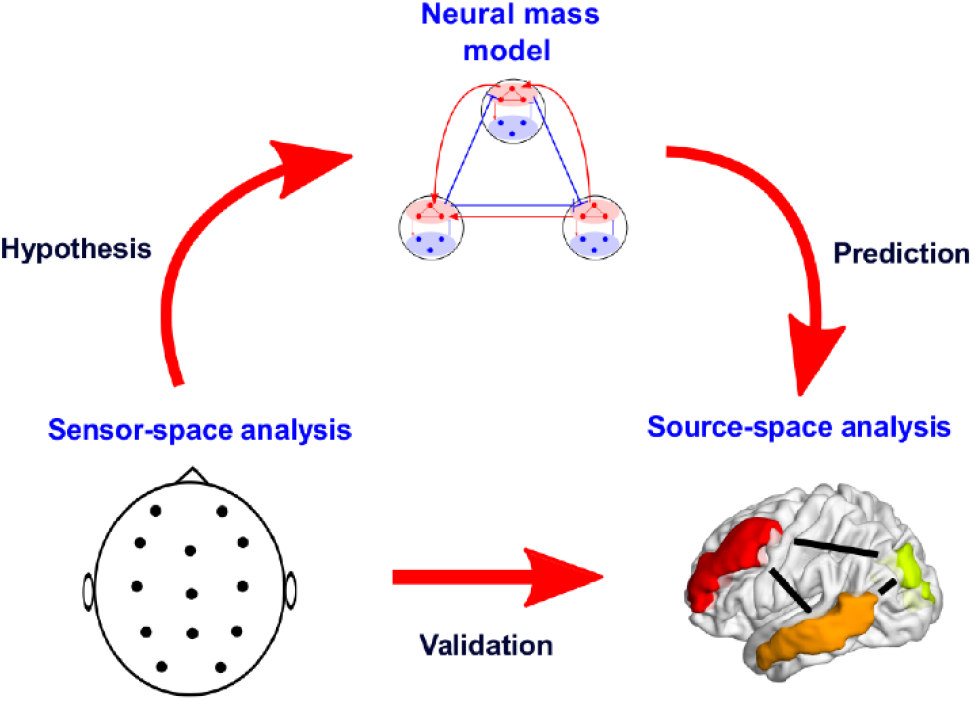
Pipeline of the study: We address an ongoing challenge of characterizing the neural mechanisms that govern the variability during multisensory speech perception. Employing the McGurk paradigm, we identify the neural signatures of the variability in the global coherence metric. We then turn to test our hypothesis – ‘dynamic and preferential coupling between sensory specific and multisensory regions underlie the observed variability’, using a neural mass model. Finally, we validate the inferences drawn from the model by analyzing pairwise coherence in the source space between concordant cortical generators of neural activity.

## Materials and Methods

### Participants

Eighteen healthy right handed volunteers (8 females, mean age 24.9, SD = 2.8) gave written informed consent under an experimental protocol approved by the Institutional Human Ethics Committee of the National Brain Research Centre, Gurgaon which is in agreement with the 2008 Declaration of Helsinki.

### Stimuli and behavioral task

A digital video of a native Hindi speaking male articulating the syllables /*pa*/, /*ka*/ and /*ta*/ was recorded and edited using – the audio editing software Audacity (www.audacityteam.org) and the video editing software Videopad Editor (www.nchsoftware.com). The duration of each video clip ranged from 1.5 to 1.7 seconds to include the neutral, mouth closed position and all the mouth movements of articulation to closing. The duration of auditory syllables in the videos ranged from 0.4 to 0.5 seconds. The stimuli (**Figure 2**) consisted of four kinds of videos: three congruent (auditory and visual matching) syllables (/*pa*/, /*ka*/, /*ta*/) and one McGurk/incongruent (auditory and visual mismatch) syllable (auditory /*pa*/ + visual /*ka*/) producing the McGurk percept of /*ta*/.

**Figure 2:**
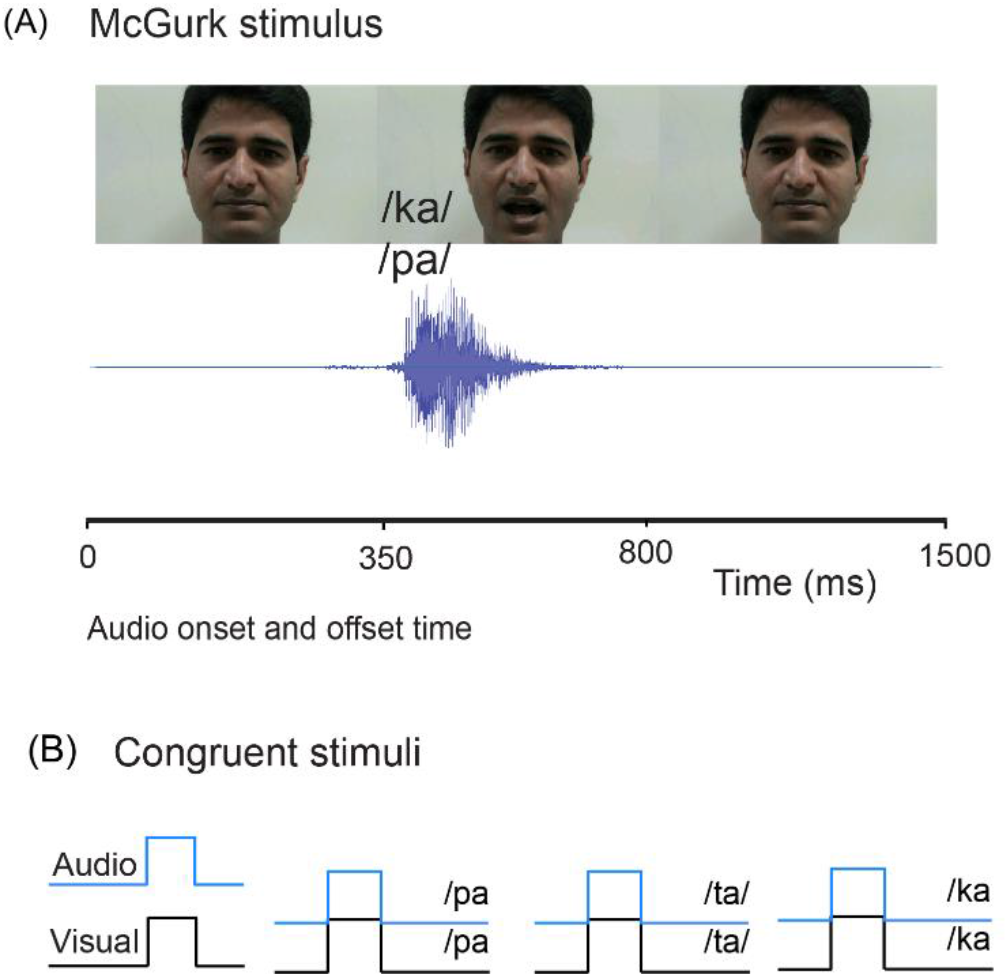
Stimuli: (A) Example trial with 3 video frames from McGurk stimulus (audio /*pa*/ + video /*ka*/) used in this experiment (top row), the audio trace of the syllable /*pa*/ presented simultaneously to the video (middle row) and the onset and offset time of the audio. (B)The congruent AV stimuli: each block represents a video with audio /*pa*/, /*ta*/ and /*ka*/ dubbed onto a video of a person articulating /*pa*/, /*ta*/ and /*ka*/ respectively.

The experiment contained five blocks – each block consisting 120 trials (30 trials of each video presented in random). Inter-stimulus intervals were pseudo-randomly varied between 1.2 seconds and 2.8 seconds to minimize expectancy effects. Using a forced choice task, on every trial, participant reported their percept by pressing a specified key on the keyboard corresponding to /*pa*/, /*ka*/, /*ta*/ or something else (others) while watching the videos.

### EEG recording and preprocessing

Continuous EEG scans were acquired using a Neuroscan system (Synamps2, Compumedics, Inc.) with 64 Ag/AgCl scalp electrodes placed on an elastic cap in a 10-20 montage. Individual electrode locations were registered using the Fastrak 3D digitizing system (Polhemus Inc.). Recordings were made against the centre (near Cz) reference electrode on the Neuroscan cap and digitized at a sampling rate of 1000 Hz. Channel impedances were monitored to be at values < 10kΩ.

Preprocessing and offline data analysis were performed using EEGLAB (Delorme & Makeig, 2004), Fieldtrip (Oostenveld et al. 2011) and custom made MATLAB scripts, Continuous data were high-passed at 0.1 Hz with finite impulse response (FIR) filters, low-passed at 80 Hz FIR and notch-filtered between 46-54 Hz to remove line noise (9^th^ order 2-pass Butterworth filter). The noisy channels were removed and interpolated, following which the data was average re-referenced. For the data analysis, epochs of 0.8s post the onset of the sound were extracted and categorized according to the type of AV stimuli - congruent AV stimuli: /*ta*/, /*pa*/, /*ka*/ and incongruent McGurk stimulus. The extracted epochs were then baseline corrected by removing the temporal mean of the EEG signal 0.2s before the onset of the sound, on an epoch-by-epoch basis. Finally, epochs with amplitudes above or below ±75µV were removed to minimize contamination from ocular (eye blinks and movements) and muscle related artifacts. To ensure comparable signal-to-noise ratio, equal number of trials were picked randomly from each participant for every kind of AV stimulus presented.

### Large-scale functional connectivity analysis

Neuronal synchronization was quantified using global coherence (Cimensar, et al. 2011). The metric quantifies the strength of covariation of neural oscillations at the global scale in specific frequency bands (Railo et al. 2011) that corresponds to specific functional state of the brain. We employed the Chronux (Mitra & Bokil, 2008) function CrossSpecMatc.m to obtain trial-wise global coherence on the 0.8s epochs sorted based on stimuli and responses. We considered 5 orthogonal discrete prolate spheroidal sequences (dpss), also known as Slepian tapers to restrict the leakage of spectral estimates into nearby frequency bins. The output variable ‘Ctot’ of the function yielded global coherence at frequency *f*.

The function initially multiplies the epochs with the specified number of Slepian tapers before performing the Fast-Fourier transform (FFT). The resulting FFT values are averaged and the cross-spectrum for all sensor combinations at frequency *f* are computed. The cross-spectral density between two sensors was computed from the tapered fourier transforms using the following equation

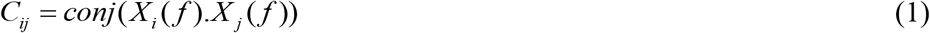

where, *C_ij_* represents the cross-spectrum, *X_i_* and *X_j_* represent the tapered Fourier transforms from the sensors *i* and *j*. Subsequently, singular value decomposition (SVD) is applied on the cross-spectral density matrix for every frequency *f* which yields the following

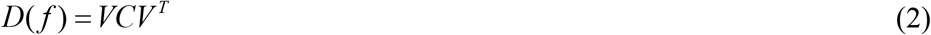

The diagonal matrix *D* comprises the values proportional to the variance explained by the orthogonal set of eigenvectors (*V*, *V^T^*). The global coherence *C_Global_*(*f*) at frequency bin *f* is subsequently computed by normalizing the largest eigenvalue (first entry of the *D*(*f*) at frequency *f* on a trial-by-trial basis for each participant employing the following equation

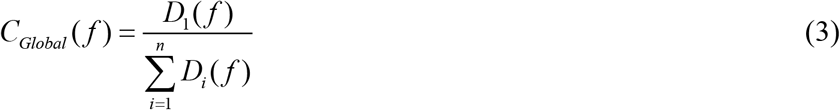

We initially analyzed if changes in global coherence values at specific frequency bands (alpha: 8-13Hz, beta: 14-30Hz, gamma: 31-45Hz) correlated with the participants’ susceptibility of McGurk perception. Participant-wise mean of the global coherence in specific frequency bands were computed and were statistically analyzed using Spearman rank correlation and t-tests.

### Architecture of large scale dynamical model of perception

To account the neuronal interactions that could facilitate changes in global coherence, we constructed a large-scale dynamical model. Striking a balance between biological realism and model parsimony, we proposed a neural-mass model comprising three nodes representing the auditory, visual and the multisensory areas (**Figure 3A**). We followed closely a previously established practice and convention in computational modeling by treating each cortical region as an individual node comprising excitatory and inhibitory population of Hindmarsh Rose (HR) neurons (Stefanescu & Jirsa, 2008). Rationally, we incorporate the following factors to reflect the biological realism in the proposed model architecture -

1. Concurring with previous evidences, auditory node was assumed to be most sensitive to ambient temporal fluctuations operating with a fast time-scale (Rosen & Howell, 2011), Visual node was assumed to be slowest in terms of sensitivity (Williams et al., 2004) and the multisensory node was assumed to operate at an intermediate time-scale.
2. Inspired by the anatomical evidences, in our model, inputs from visual node were directed to the auditory cortex in two ways: i) direct influence in a feedforward manner (Falchier et al., 2002; Rockland & Ojima, 2003; Wallace et al., 2004) and ii) in feedback manner via the multisensory association node (Bizley & King, 2012).
3. Excitatory and inhibitory afferents on the spatially extended and geometrically aligned pyramidal neurons result in post-synaptic potentials (Kirschstein & Köhling, 2009), These communication emerge as EEG patterns. Hence, we used a population of excitatory and inhibitory neurons in each node (**Figure 3B**). Following the approach used by Stefanescu and Jirsa (Stefanescu & Jirsa, 2008), we employed a total of 150 excitatory (*N_E_* = 150) and 50 inhibitory (*N_I_* = 50) HR neurons, maintaining a ratio of 3:1. Importantly, inhibitory neurons within an individual node were not coupled directly with other inhibitory neurons based on the logic that these connections in microcircuits are typically sparse in nature (Wilson & Cowan, 1972; Stefanescu & Jirsa, 2008).

**Figure 3:**
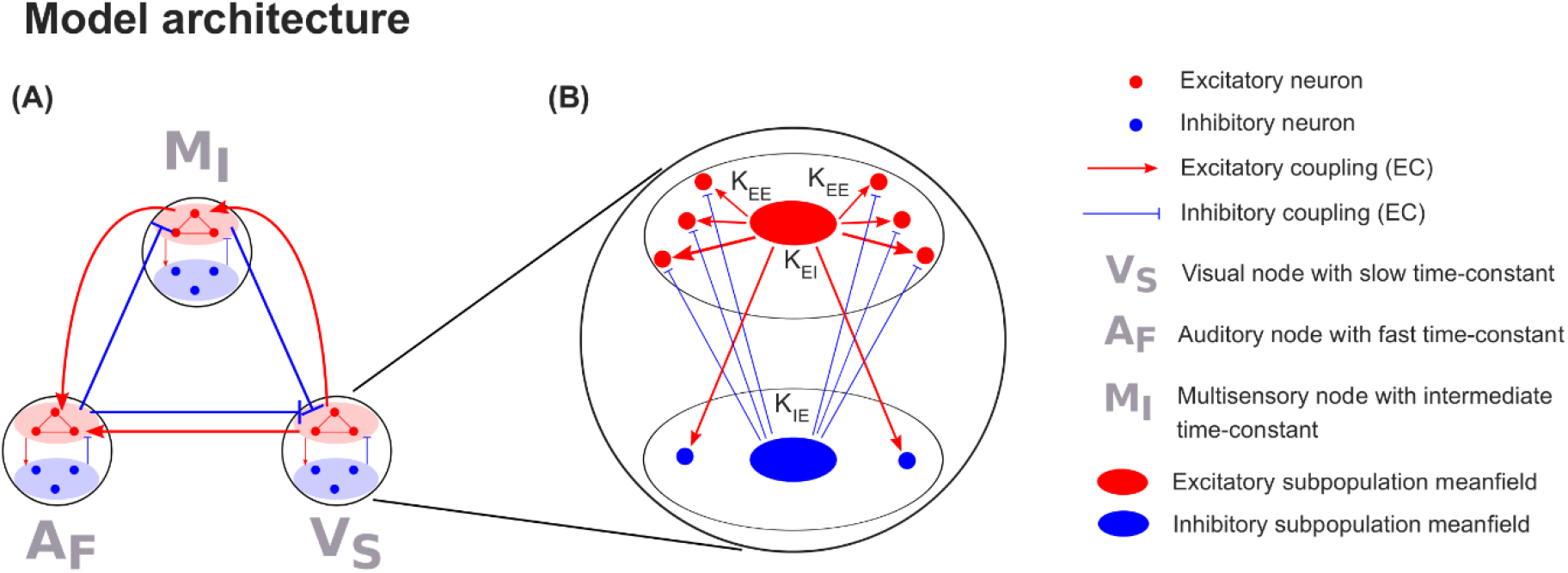
Model architecture: (A) Representation of the model comprising three nodes signifying auditory (fast time-constant), visual (slow time-constant) and higher order multisensory regions (intermediate time-constant). Excitatory influences are balanced by a reciprocal inhibitory connection. (B) Representation of each node comprising of network of 150 Hindmarsh-Rose excitatory and 50 inhibitory neurons. Each neuron can exhibit isolated spiking, periodic spiking and bursting behavior. The detailed model parameters are presented in **Table 1**.

Incorporating the aforementioned factors, we defined a neural mass model comprising three sets of ordinary differential equations for each excitatory and inhibitory HR neurons. The three variables 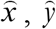 and 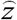 account for the membrane potential dynamics, fast and slow gating currents respectively. Thus, the entire network can be represented as a network of coupled non-linear differential equations containing -

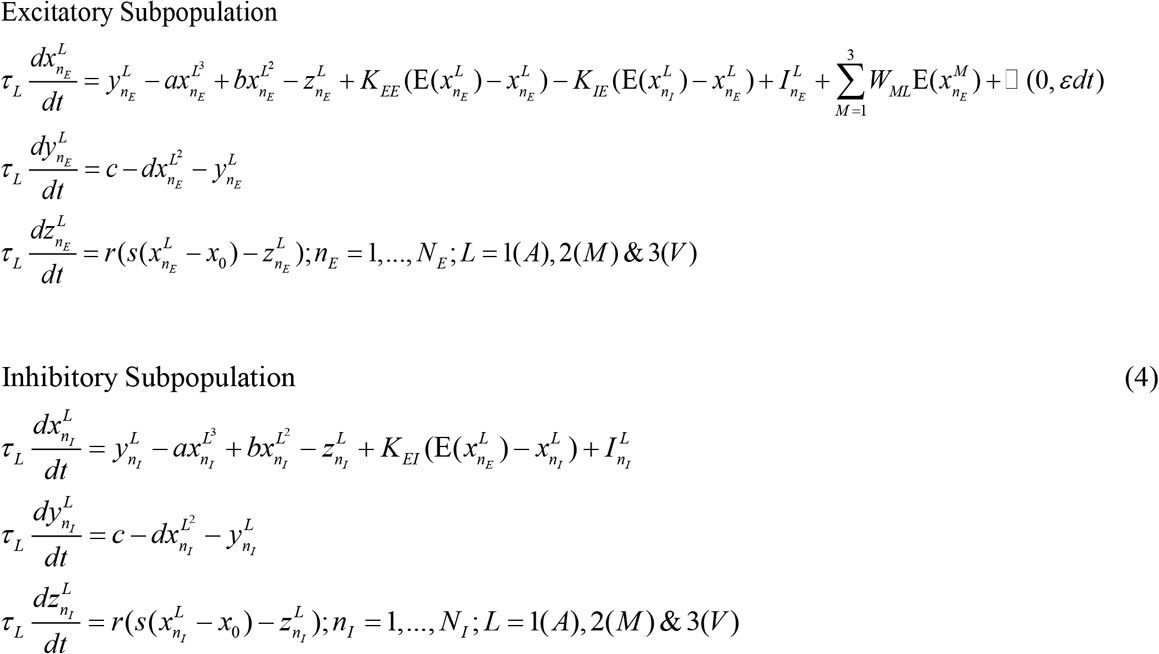

where *L*: A, V and M represent auditory, visual and multisensory nodes respectively. The nodes were driven by a noise of normal distribution □ (0,*εdt*), where *ε* is 1*e* −1. Auditory node had the fastest (*τ_A_* ~ 0.05*ms*) and visual node had the slowest time-constant (*τ*_*V*_ ~ 2.5*ms*). Time-constant of the multisensory node was chosen to be in between the two (*τ_M_* ~ 1*ms*) as it integrates information from both the modalities. The mean activity of excitatory neurons in a given node (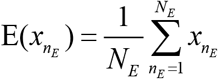) influencing the neuronal activity of other nodes was governed by coupling parameters: *W_AV_* (auditory-visual coupling), *W_AM_* (auditory-multisensory coupling) and *W_VM_* (visual-multisensory coupling). Positive and negative values of the coupling parameters reflect excitatory and inhibitory coupling respectively. Inhibitory couplings were chosen to maintain a balance with excitation. For instance, visual node’s excitatory coupling of +*W_AV_* on auditory node was balanced with inhibitory coupling of the same strength (−*W_AV_*) from the auditory node. Also, in such a configuration, visual node is being the source of excitation, is referred as the source whereas auditory node is referred as the sink since all excitatory couplings are directed towards it. The multisensory node in such configuration behaves as both source and sink.

We place each individual neuron in a dynamical regime where both spiking and bursting behavior is possible depending on the external input current (I) that enters the neuron when other parameters are held constant at the following values:

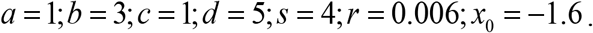

(Stefanescu & Jirsa, 2008).

The coupling between neurons within a node was linear and its strength was governed by the following parameters: *K_EE_* for excitatory-excitatory coupling, *K_EI_* for excitatory-inhibitory coupling and *K_IE_* for inhibitory-excitatory coupling. Within a node, the relation between *K_EE_* and *K_IE_* was captured by the ratio 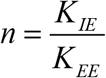. Electrophysiological evidences indicate enhanced oscillatory power in delta (1-4 Hz) and alpha (8-12 Hz) bands during rest (Gold et al., 2006). To emulate the characteristics of the large-scale network during rest, we chose the value of the ratio, *n* = 3.39 where the average oscillatory activity of the nodes in a disconnected network i.e. in the absence of stimulus (*μ*(*I*_*A*,*V*, *M*_) = 0.1; baseline) was higher in delta and alpha bands.

The external currents to both the excitatory and inhibitory subpopulations were drawn from a Gaussian distribution where *μ* and *σ* represent the mean and standard-deviation. As the input stimulus relays to auditory, visual and multisensory regions via thalamus, we interpret lateral geniculate nucleus (LGN) and medial geniculate nucleus (MGN) to be the source of external current (*I_A_*, *I_V_* and *I_M_*) – a pulse of 450 ms in the nodes when the model was simulated for 1 second. In rhesus monkey, the projections of MGN to multisensory areas like STS were found to be sparse (Yeterian & Pandya, 1989). Therefore, we choose lower mean value of external current to the multisensory node (*μ*(*I_M_*) = 0.85) in comparison to visual (*μ*(*I_V_*) = 2.8) and auditory (*μ*(*I_V_*) = 2.8) node. All parameters are presented together in **Table 1**.

**Table 1:** Description of the model parameters and their corresponding values used in the model.

| Parameter | Value | Description of parameters |
| --- | --- | --- |
| $a, b, c, d$ | 1, 3, 1, 5 | Constants for faster variables x & y |
| $r$ | 0.006 | Lower value of r is responsible for slower time-scale of z |
| $s$ | 4 | Constant affecting slower variable z |
| $x_0$ | -1.6 | $(x_0, y_0, 0)$ is a stable equilibrium point of Hindmarsh-Rose model. |
| $N_E, N_I$ | 150, 50 | Number of excitatory and inhibitory neurons in each node |
| $\mu, \sigma$ | 2.8, 0.4 | Mean and dispersion of input current in each node |
| $I_{n_E}, I_{n_I}$ | Computed from $\mu$ and $\sigma$ | Input current for excitatory and inhibitory neurons in each node |
| $\tau_A, \tau_V, \tau_M$ | 0.05, 2.5, 1<br>in milliseconds | Time-scales for auditory, visual and multisensory nodes |
| $\varepsilon$ | 0.1 | Constant affecting the dispersion of noise |
| Within nodes connectivity |  |  |
| $K_{EE}$ | 0.5 | Coupling between excitatory neurons |
| $K_{EI}$ | 0.5 | Excitatory to inhibitory coupling |
| $n$ | 3.39 | $K_{IE}$ to $K_{EE}$ ratio |
| $K_{IE}$ | Computed from n and $K_{EE}$ | Inhibitory to excitatory coupling |
| Between nodes connectivity |  |  |
| $W_{AV}, W_{AM}, W_{VM}$ | Explored in the range 0 to 1 (or -1 to 0) | |
| State Variables |  |  |
| $x_{n_E}, x_{n_I}$ | Membrane potentials for excitatory and inhibitory subpopulation | |
| $y_{n_E}, y_{n_I}$ | Spiking variables for excitatory and inhibitory subpopulation | |
| $z_{n_E}, z_{n_I}$ | Bursting variables for excitatory and inhibitory subpopulation | |

We hypothesized that variability in global coherence patterns to emerge from differences in the coupling between the sensory specific and multisensory brain regions operating at fast, slow and intermediate time scales respectively. Therefore, we sought to explore the global coherence while varying the coupling strength among the nodes in our model.

In summary, we considered different architectures or connectivity motifs, under the constraint that any reciprocal coupling comprised a forward excitatory set of connections and a reciprocal (and balanced) inhibitory set of connections. In what follows, we will designate the source of the excitatory connections as a source – and the source of recurrent inhibitory connections as a target.

### Source-space connectivity

To validate the inferences gathered from our model, we turned to the empirical data again and undertook source-space connectivity analysis (see **Figure 1**). We employed dynamic imaging of coherent sources (DICS) (Gross et al., 2001)- a frequency domain adaptive spatial filtering algorithm - to identify the sources of global coherence in specific frequency bands. This algorithm has proven to be particularly powerful in localizing oscillatory sources of cortical activity with well-established hypothesis (Halder et al., 2019). Customized MATLAB codes employing Fieldtrip toolbox were used. Using ft_prepare_leadfield.m and employing the Boundary Element Method (BEM), we generated the leadfield matrix for the Colin27 brain template. The leadfield matrix corresponds to the tissue and geometrical properties of the brain represented as discrete grids or voxels. Then, we employed ft_freqanalysis.m to evaluate the crossspectral matrices of the epochs pooled over all the participants. Leadfield matrix and the cross-spectral matrices were used to compute the common spatial filters that regulate the amplitude of brain electrical activity passing from a specific location while attenuating activity originating at other locations. The distribution of the output amplitude of the spatial filters provides the metric for source localization.

To reduce the dimensionality in our source-space connectivity analysis, we adopted a parcellation scheme before generating source-time series. We combined the source estimates of the voxels using the labeling according to Automated Anatomical Labelling (AAL) atlas (Tzourio-Mazoyer et al., 2002). Then, for reconstructing the parcel-wise time series, we projected single trial epoched time series to the source-space by multiplying them with spatial filter of the respective parcel obtained employing DICS. Subsequently, we computed the coherence between relevant parcels for rare and frequent perceivers explicitly.

Since the creation of common spatial filter required multiple trials and participants to reconstruct one reliable estimate of source time series, we resort to bootstrapping to compare the differences in pairwise source-level coherence between rare and frequent perceivers. Initially for each relevant parcel, from the trials pooled over participants in each group (rare and frequent), 500 trials were randomly picked to form two sets of random partition. Subsequently, pairwise coherence between the parcels were computed on a trial-by-trial basis and averaged for rare and frequent perceivers separately.

The aforementioned steps were repeated 500 times to generate a distribution of pairwise coherence values for frequent and rare perceivers. Significance of pairwise coherence difference values between the rare and frequent perceivers were evaluated from bootstrapped estimates.

### Data & Code Availability

Raw and processed data used for generation of figures from empirical data as well as large-scale dynamical model will be shared on request to Corresponding author AB.

## Results

### Variability in multisensory perception

Incongruent McGurk stimulus, visual /*ka*/ paired with audio /*pa*/ also referred as /*pa*/*-*/*ka*/, and three kinds of congruent AV stimuli - /*ta*/−/*ta*/, /*pa*/−/*pa*/, /*ka*/−/*ka*/ were presented to all participants. The sequence of presentation was randomized. As the participants observed each of the stimuli, they were instructed to report with a set of four keys if they had heard /*ta*/, /*pa*/, /*ka*/ or ‘something else’.

Response tendency (*inter-trial variability*), computed as the relative proportion of illusory (/*ta*/) responses in all McGurk trials across all participants, showed that /*ta*/ percept was reported in 58.19% (SD = 32.33) of trials, whereas unisensory /*pa*/ percept was reported in 37.56% (SD = 31.79) trials (Figure 4A). However, paired two-sample *t-tests* comparing the percentage of illusory /*ta*/ and unisensory /*pa*/ percept over all the participants revealed no significant difference (*t_17_*= 1.36, *p* = 0.19). Congruent AV stimuli (/*ta*/, /*pa*/ and /*ka*/) were correctly identified in 96.56% (SD = 2.88) trials by all the participants.

**Figure 4:**
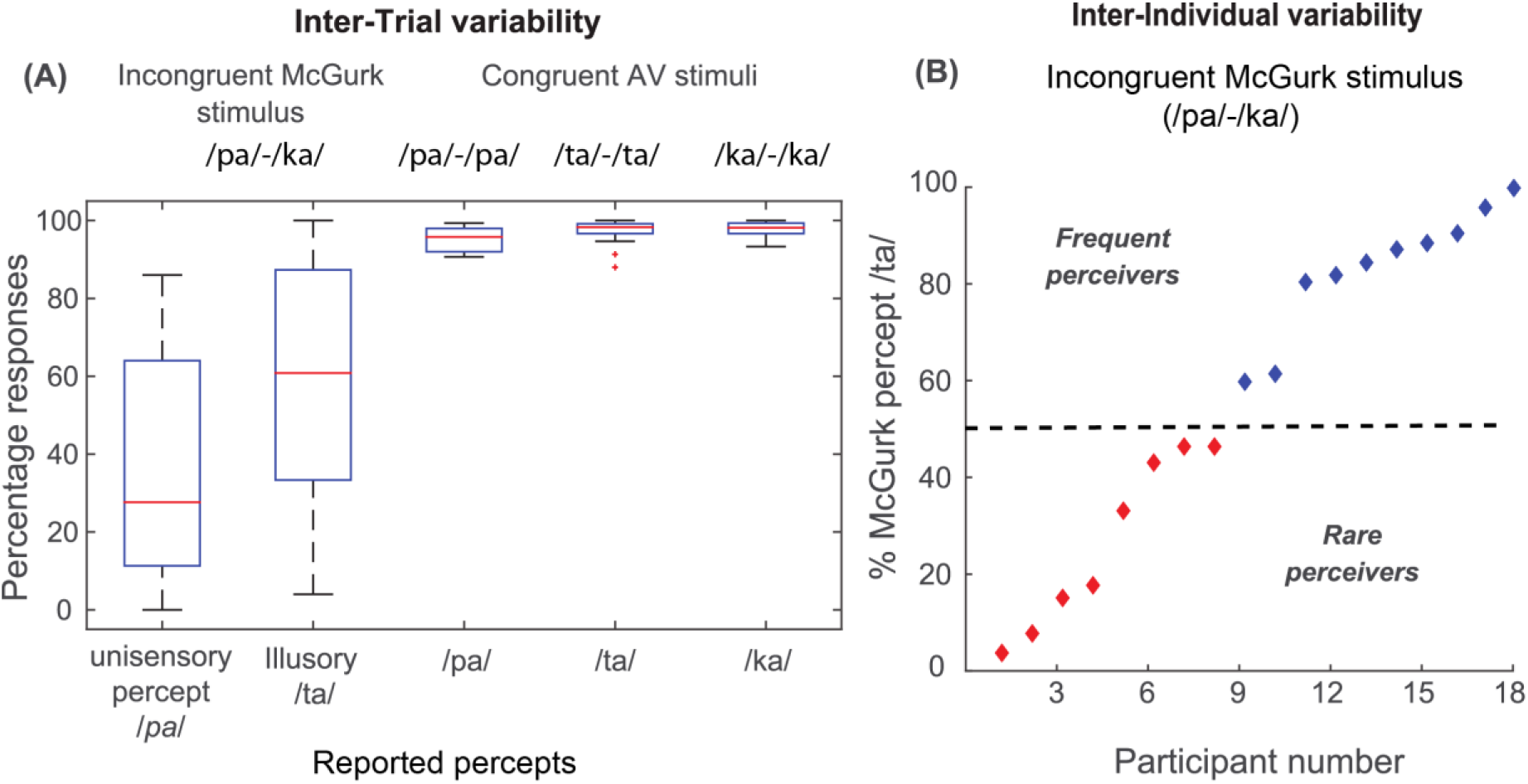
Behavior: Behavior: (A) *Inter-trial variability* - Percentage of /*ta*/ (illusory) and /*pa*/ (unisensory) percept during the presentation of McGurk stimulus and the hit rate during the presentation of congruent AV stimuli (/*pa*/, /*ta*/ and /*ka*/) averaged over all the participants. The red mark in each box plot indicates the median (B) *Inter-individual variability* - Propensity of McGurk effect for each of the 18 participants expressed as the percentage of /*ta*/ percept during the presentation of McGurk stimulus. The participants were categorized as frequent perceivers (blue diamonds, >50%) and rare perceivers (red diamonds, <=50%).

A high degree of *inter-individual* variability in McGurk susceptibility was observed (**Figure 4B**) while analyzing participant-wise /*ta*/ percept during McGurk stimulus. The participants were classified based on their McGurk susceptibility in the incongruent trials into two groups – rare perceivers (<50% /*ta*/ percept) and frequent perceivers (>50% /*ta*/ percept). Eight of the eighteen participants elicited <50% propensity of perceiving the McGurk effect. A two-way analysis of variance with congruent AV stimuli (/*pa*/, /*ta*/ and /*ka*/) and participant groups (frequent and rare) as factors revealed that only the type of congruent stimulus affect the hit rate, *F*(2,53) = 4.44, *p* = 0.02, and the effect of participant group (frequent or rare) on the hit rate was not significant, *F*(1,53) = 4.18, *p* = 0.05. Also, there was no evidence of interaction between the congruent AV stimulus type and the participant group *F*(2,53) = 0.07, *p* = 0.937. We further performed paired two-sample *t-test* to compare the differences in the hit rate during the congruent stimuli. We observed that hit rate during congruent AV stimulus /*pa*/ was significantly lower that hit rate during congruent AV stimuli: /*ta*/ (*t*_17_ = −2.06, *p* = 0.047) and /*ka*/ (*t*_17_ = −2.97, *p* = 0.005).

### Large-scale functional connectivity in EEG sensor space

The emergence of specific cognitive trait is subserved by the coordination of distributed functionally specialized neuronal assemblies or brain regions (Bressler, 1995; Varela et al., 2001; Bressler & Menon, 2010). Based on our previous study (Kumar et al., 2016), we hypothesized that neural signatures of inter-individual can be quantified using the global coherence metric that capture the degree of neuronal synchronization over the entire brain from EEG scalp-level recordings.

We observed a significant negative correlation between each participant’s alpha band global coherence and their McGurk susceptibility during incongruent McGurk condition (/*pa*/−/*ka*/), (*r_s_* (18) =−0.49, *p=*0.04) (**Figure 5A**), congruent /*ta*/−/t*a*/ (*r_s_* (18) =−0.51, *p=*0.03) (**Figure 5B**) and /*ka*/−/*ka*/ (*r_s_* (18) =−0.49, *p*=0.04) (**Figure 5C**) stimuli. However, alpha modulation among participants during congruent /*pa*/−/*pa*/ condition were not correlated with their McGurk susceptibility (*r_s_* (18) =−0.18, *p=*0.47) (**Figure 5D**). Evidently, there were also no correlation between global coherence in any other frequency bands and McGurk susceptibility during incongruent McGurk (/*pa*/−/*ka*/), theta: *r_s_* (18) =−0.29, *p*=0.22., beta: *r_s_* (18) =−0.21, *p*=0.39, gamma: *r_s_* (18) =−0.10, *p* = 0.45; congruent /*pa*/, theta: *r_s_* (18) =−0.38, *p*=0.12, beta: *r_s_* (18) =−0.14, *p*=0.57, gamma: *r_s_* (18) =−0.15, *p*=0.53; congruent /*ta*/ theta: *r_s_* (18) =−0.36, *p*=0.14, beta: *r_s_* (18) =−0.17, *p*=0.50, gamma: *r_s_* (18) =−0.19, *p*=0.42, and congruent /*ka*/ (theta: *r_s_* (18) =−0.31, *p*=0.20, beta: *r_s_* (18) =−0.22, *p*=0.37, gamma: *r_s_* (18) = −0.16, *p*=0.50 stimuli.

**Figure 5:**
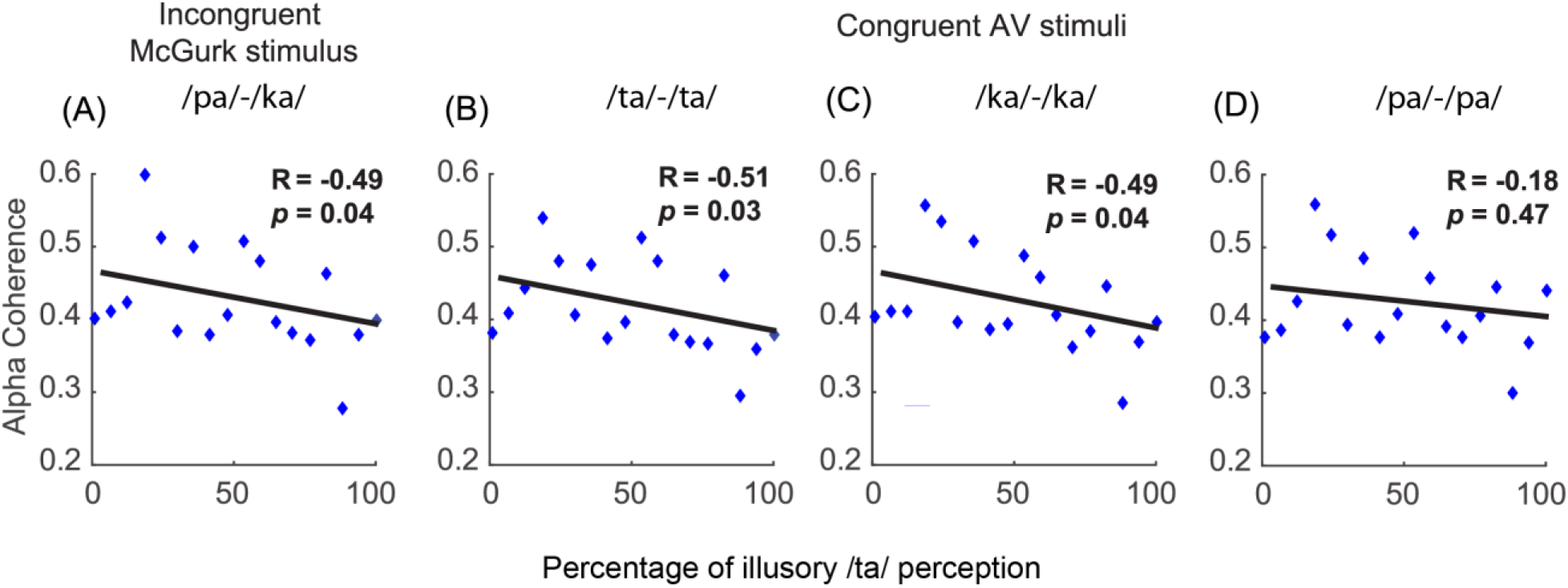
Large scale functional connectivity: Mean alpha band coherence in each participant plotted against that participant’s susceptibility of McGurk effect during stimulus: (A) McGurk (B) Congruent /*ta*/ (C) Congruent /*ka*/ (D) Congruent /*pa*/

### A simple interacting neural mass model with fast, intermediate and slow time-constants explains desynchronization of alpha band global coherence

Global coherence dynamics may reflect the potential operation of a large-scale network that support multisensory speech perception. We hypothesized that the variability in McGurk percept across participants to be grounded in specific interactions between neural assemblies (**Figure 6**). In order to account for the observed global coherence patterns, we employed a neural mass model comprising three nodes broadly representing the auditory and visual areas in the sensory cortices, and the multisensory node representing the collection of multisensory (or modality-nonspecific) areas. Each node comprised a population of excitatory and inhibitory Hindmarsh-Rose (HR) neurons (Hindmarsh & Rose, 1984) operating at different physiologically motivated time-scales (see Materials and Methods for more details). Model parameter details are presented in **Table 1**.

**Figure 6:**
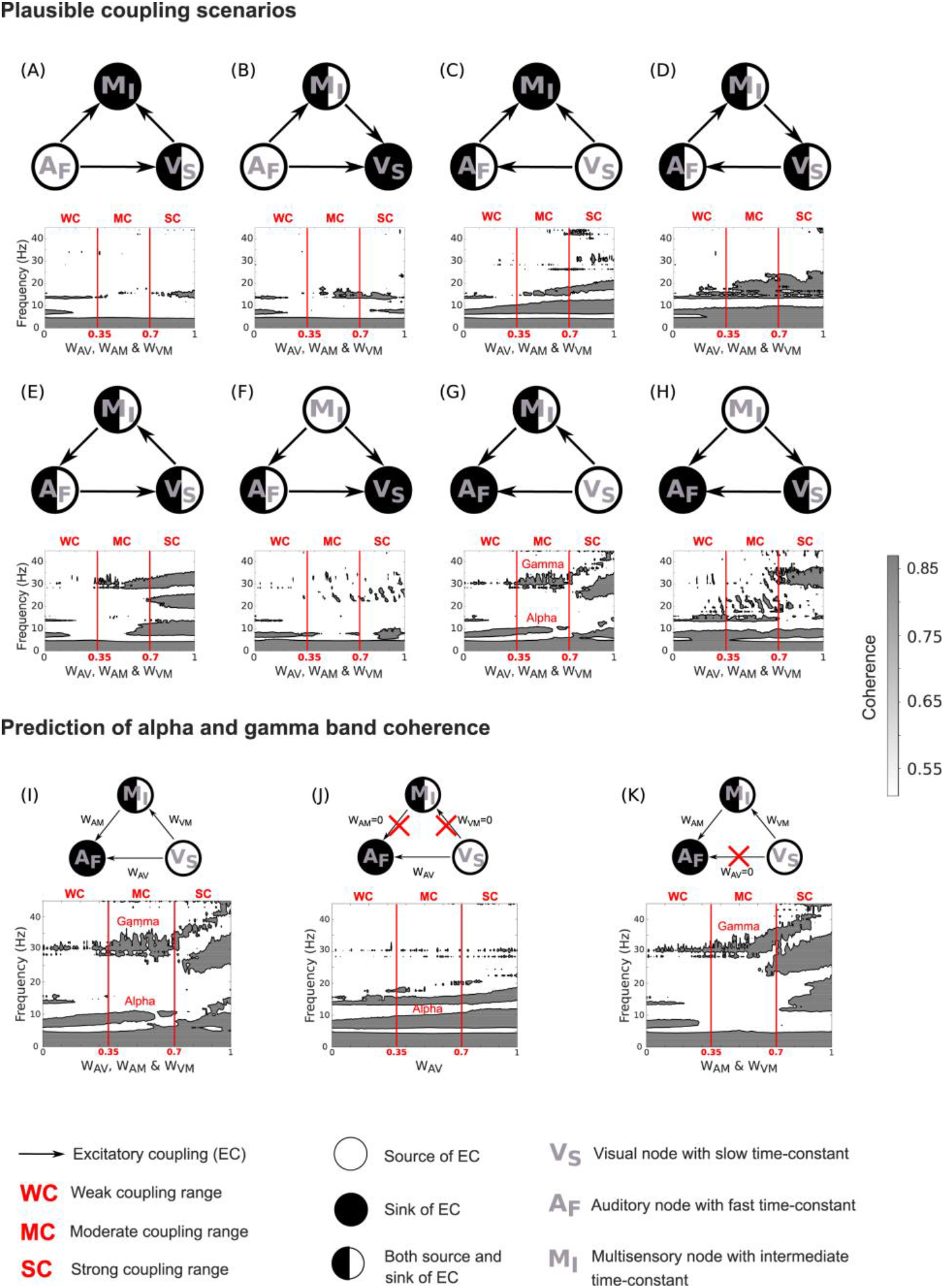
Prediction of alpha and gamma coherences from neural mass model: (A) - (H) Exploration of various coupling scenarios to identify if it is possible to generate alpha and gamma coherence in moderate coupling range. Only the model configuration (G) exhibits concurrent coherence in the alpha and gamma band in the moderate coupling range. I) Alpha and gamma band coherences co-exist in moderate coupling range when visual node is source and auditory node is the sink J) Only direct A-V coupling generates alpha coherence independently. K) Indirect A-V coupling via multisensory node generates gamma coherence at the limit case scenario of weak direct coupling.

We sought to explore the global coherence while varying the configuration and strength of coupling among the nodes in our model. For instance, in **Figure 6A** auditory node’s excitatory coupling of +*W_AV_* on multisensory node was balanced with inhibitory coupling of the same strength (−*W_AV_*) from the multisensory node. In such a configuration, auditory node is being the source of excitation, is referred as the source whereas multisensory node is referred as the sink since all excitatory couplings are directed towards it. The visual node in such configuration behaves as both source and sink.

The co-existence of coherence in alpha and gamma band facilitated by the two time scales of visual (slow) and auditory (fast) node was observed in network configuration wherein visual node operates as the source and auditory node acts as a sink (**Figure 6G**). Moreover, the concurrence of the coherence in alpha and gamma bands emerged in the coupling between auditory and visual regions (coupling parameter ranging between 0.35 and 0.7) via two routes – 1) direct coupling (*W_AV_*) and 2) indirect coupling (*W_AM_* & *W_VM_*) involving multisensory node (M). We assumed coupling strength less than 0.35 to be weak coupling (WC), coupling strength between 0.35 and 0.7 to be moderate coupling (MC) and coupling greater than 0.7 to be strong coupling (SC). In SC range, we observed high coherence in alpha band and gamma bands, however, no peak was observed in alpha band. In the MC range, none of the model configurations were able to generate concurrent coherent states in alpha and gamma bands (**Figure 6A-H**), except **Figure 6G**, where visual node acts as the source of excitation. Furthermore, direct coupling between auditory and visual node alone (*W_AV_* ranges from 0 to 1, *W_AM_* & *W_VM_* = 0) results only in alpha band coherence (**Figure 6J**). Interestingly, emergence of only gamma band coherence was observed when the indirect coupling in the model were varied between A-V nodes via multisensory node (*W_AM_* & *W_VM_* range from 0 to 1, *W_AV_* = 0) in MC range (**Figure 6K**).

#### Direct Audio-Visual interaction underpins Inter-Individual Variability

Our empirical findings indicate a significant negative correlation between participant-wise alpha band coherence and McGurk susceptibility. We hypothesized that mediation of interaction between auditory (A) and visual (V) node via the multisensory node (M) (i.e. stronger A-M and V-M coupling than A-V coupling) to be characteristic of participants with higher susceptibility to the McGurk effect. To test this hypothesis, a balanced network coupling state, *W_AM_* = *W_VM_* = *W_AV_* = 0.35 where alpha and gamma band coherences co-exist was considered, following which the change in the coherence peaks with decreasing direct A-V coupling (*W_AV_*) was studied. A suppression of alpha coherence peak with decrease in *W_AV_* was observed (**Figure 7**); however, gamma coherence peak remained more or less unchanged. Only when A-V coupling in the computational model became negligible (*W_AV_* < 0.05), disappearance of alpha coherence peak was observed. We use this basic observation to revisit the empirical data and ask whether there was evidence for a disconnection or uncoupling between auditory and visual sources in subjects who evinced differential alpha coherence.

**Figure 7:**
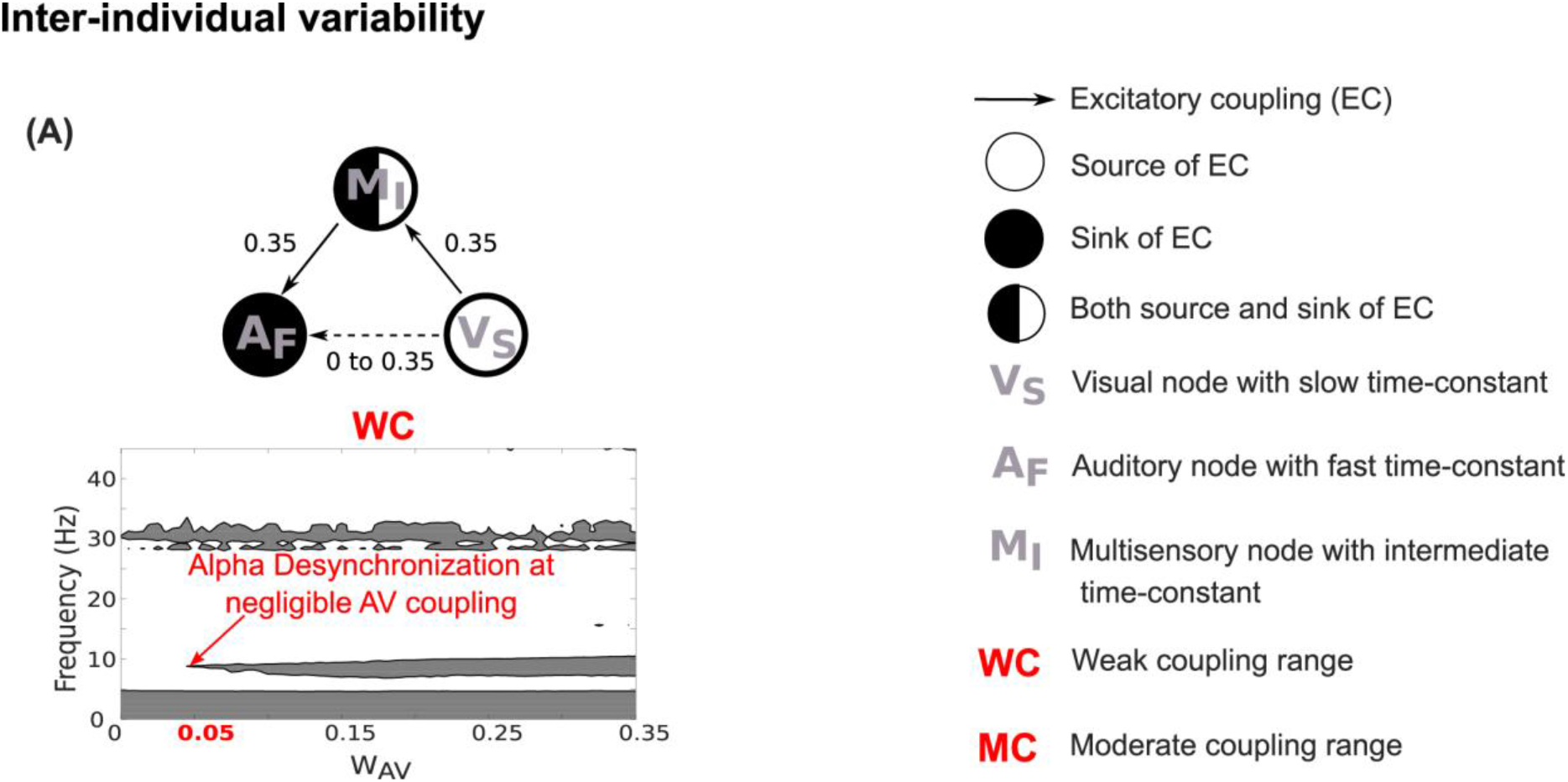
Mechanistic understanding of Inter-individual variability: A) Alpha de-synchronization characteristic of frequent perceivers resulted due to negligible A-V coupling. **Note**: (reciprocal) inhibitory connections have been omitted for clarity.

### Validation of model predictions using source-connectivity analysis

Finally, we obtained source level time series for each brain parcel by projecting single trial time series to the source-space by multiplying them with spatial filter of the respective parcel using voxel to parcel mapping provided by AAL atlas (see materials and methods for details). Subsequently, the pairwise coherence - a measure of functional connectedness, between the parcels associated with auditory and visual areas were computed. Auditory specific parcel – Superior Temporal Gyrus (STG) and the visual parcels – Superior Occipital Cortex (SOC), Medial Occipital Cortex (MOC) and Inferior Occipital Cortex (IOC) were identified. A significant increase of 0.8% and 0.5% in the functional connectivity in rare perceivers than frequent perceivers was observed between left STG - left SOC (mean coherence increase = 0.0071, SD = 0.0015), and between left STG – left MOC (mean coherence increase = 0.0053, SD = 0.0020) respectively (Figure 8, Table 2). Percentage increase in the aforementioned coherence differences in rare perceivers compared to frequent perceivers between the auditory and visual parcels in both hemispheres are summarized in the Table 2. Achieved significance levels were estimated using normal intervals as the bootstrapped estimates of coherence differences followed normal distribution (Wasserman, 2005). Normality of each coherence distribution were tested using Lilliefors test where the null hypothesis that bootstrapped coherence values comes from a normal distribution couldn’t be rejected at 95% significance. To cross-check the results we ran several permutations of the comparison, and the main pattern of left STG-SOC and left STG –MOC connectivity being significantly higher in rare perceivers did not change.

**Figure 8:**
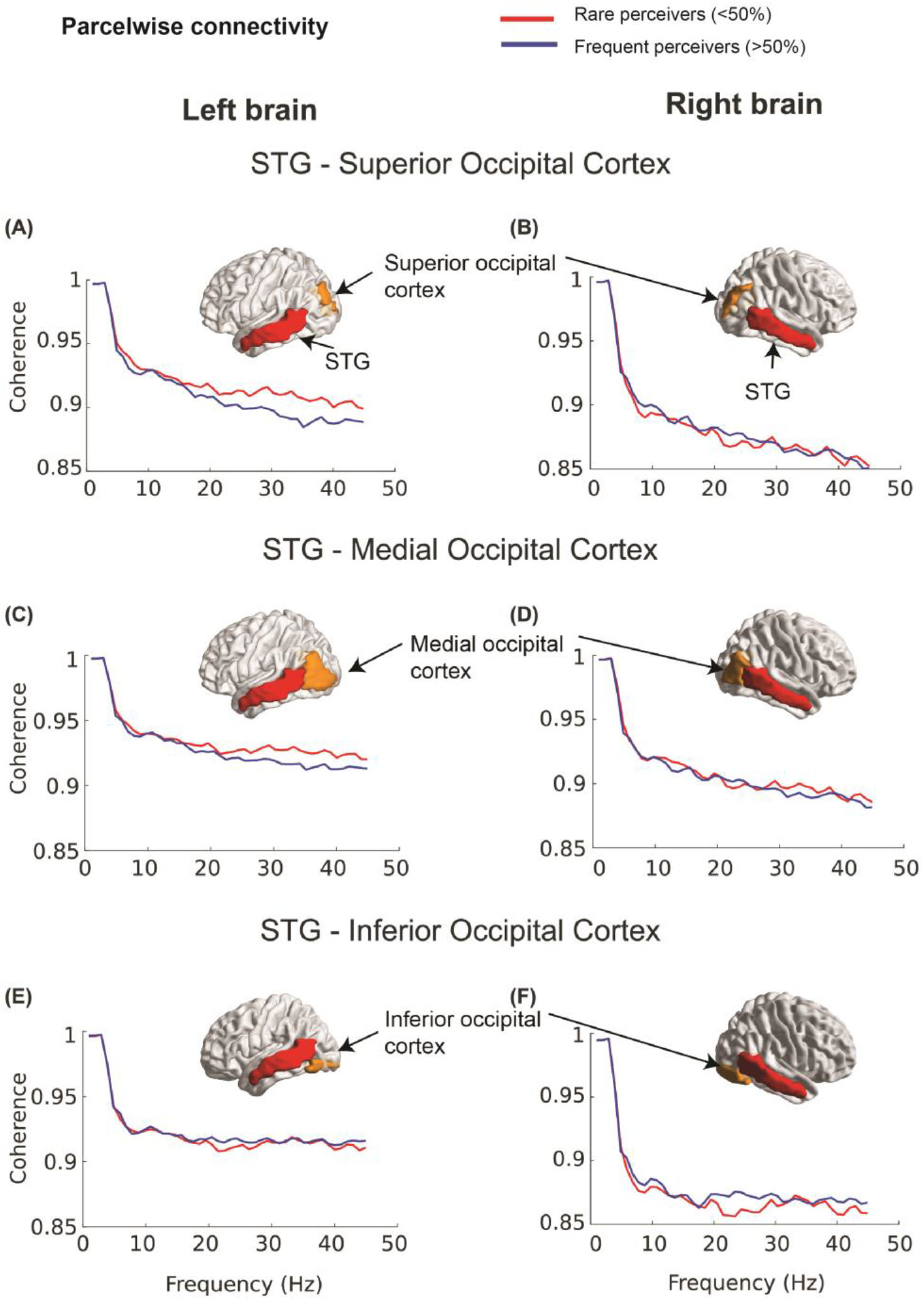
Pairwise connectivity between the parcels: Pairwise coherence between: (A) left Superior Temporal Gyrus (STG) and left Superior Occipital Cortex (SOC) (B) right STG and right SOC (C) left STG and left Medial Occipital Cortex (MOC) (D) right STG and right MOC (E) left STG and left Inferior Occipital Cortex (IOC) (F) right STG and right IOC in rare and frequent perceivers The brain images represent parcels between which the pairwise coherence is computed.

**Table 2:** Summary of the source-level coherence difference between the parcels from rare and frequent perceivers (bootstrapped estimates, see Methods).

| *Connectivity* | *Left hemisphere* |  |  |  | *Right hemisphere* |  |  |  |
| --- | --- | --- | --- | --- | --- | --- | --- | --- |
|  | %<br>increase | Mean<br>difference | SD | Achieved<br>significance<br>level (ASL)<br>*p* | %<br>increase | Mean<br>difference | SD | Achieved<br>significance<br>level (ASL)<br>*p* |
| *STG-SOC* | 0.8 | 0.0071 | 0.0015 | <0.0001 | 0.2 | 0.0016 | 0.0017 | 0.53 |
| *STG-MOC* | 0.5 | 0.0053 | 0.0020 | <0.0001 | 0.4 | 0.0038 | 0.0015 | 0.12 |
| *STG-IOC* | 0.3 | 0.0035 | 0.0010 | 0.16 | 0.3 | 0.0028 | 0.0016 | 0.09 |

## Discussion

An ongoing challenge in multisensory speech perception community is to accurately identify and characterize the neural mechanisms that govern the perceptual variability across individuals. Traditionally, studies have focused on identifying the neural correlates in terms of candidate brain regions or interactions among specific regions of interest that are responsible for the observed heterogeneity (Nath & Beauchamp, 2012). A more emerging understanding however, suggest the existence of large-scale networks of brain regions facilitating perceptual processing (Keil et al 2012; Beauchamp, 2015; Kumar et al 2017). Neuronal oscillations has been identified as a key marker of neuronal information processing in the relevant brain regions that needs to be fully explored to shape an individual’s perceptual experience (Keil et al 2012; Kumar et al 2016, 2017; Keil & Senkowski, 2018). Robust oscillations observed from macroscopic recordings such as EEG/ MEG are an outcome of network interactions among local subpopulations of excitatory and inhibitory neurons (Wilson & Cowan, 1972; Deco et al., 2010; Becker et al., 2015). Empirically such interactions result in global coherence dynamics (Kumar et al., 2017). Our study demonstrates how distinct global coherence patterns correlate with inter-individual heterogeneity. Subsequently, using a computational model of interacting large-scale brain networks, we explain the plausible neural mechanisms that generate the coherence dynamics associated with the observed heterogeneity. Finally, we validate the plausible mechanism that emerged from the model about the connection parameters among interacting neural-subsystems using source-space functional connectivity analysis. Put together, we present a biologically plausible mechanistic proposal that underlie the observed inter-individual variability in multisensory speech perception.

The key observations in our study are – 1) susceptibility of individuals’ McGurk effect is negatively correlated to their alpha band global coherence - indicating that the degree of synchronization of large-scale neural assemblies in the alpha band underlie inter-individual variability 2) weakened direct A-V coupling in the neural mass model leads to the decrease in alpha band coherence and 3) individuals with higher McGurk susceptibility elicited decreased pairwise connectivity between the sensory specific (audio and visual) areas than individuals with lower McGurk susceptibility.

### Global coherence as the index of perceptual variability

Inter-individual differences observed in our study replicate (including reports from our group) and extend previous studies. 44% of our participants had more than 80% McGurk percept which lies within the range of 26% to 100% reported in previous studies (McGurk & Macdonald, 1976; MacDonald, et al., 2000; Benoit, et al., 2009; Nath & Beauchamp, 2012) (**Figure 4A**). Also, our mean McGurk percept (collapsed across participants) of 58% is within the literature range of 24% to 100% (McGurk & Macdonald, 1976; MacDonald, et al., 2000; Benoit, et al., 2009; Nath & Beauchamp, 2012; Strand et al., 2014) (**Figure 4A, B**). Hence, the behavioral results give confidence that the corresponding brain patterns will be of robust significance and general applicability. Our results show that the extent of de-synchronization of the cortical network in the alpha band indexes an individual’s McGurk susceptibility (**Figure 5A, B, C**). This finding has two key elements. First, the inverse relationship between McGurk susceptibility and alpha band global coherence was consistent for McGurk stimulus and the congruent AV stimuli (/*ta*/ and /*ka*/) as well. This highlights that our observations indeed reflect inter-individual differences/ perceptual categorization and not necessarily stimulus configurations. Second, it pinpoints that inter-individual variability is contingent on the functional state of the brain which can be captured by exploring the dynamics of the large-scale cortical network. The syllables /*ta*/ and /*ka*/ can be difficult to distinguish with visual cues alone, but acoustically are different. Therefore, coincidence of AV cues allows both the group of perceivers to reap the benefits of AV integration during congruent AV stimuli /*ta*/ and /*ka*/. However, the bilabial syllable /*pa*/ is very different visually from /*ta*/ and /*ka*/. Hence, identification of the congruent AV syllable /*pa*/ does not necessarily require AV integration. Therefore, we do not see any significant correlation during congruent /*pa*/ stimulus (**Figure 5D**).

Previous evidences accentuate the modulations in alpha band coherence to central executive processes (Klimesch, 1999; Sauseng et al., 2005), that are postulated to be involved in allocating working memory storage to phonological loop that maintains verbal information, and the visuo-spatial sketchpad that maintains transient visuo-spatial information (Baddeley, 1992). Such processes are involved in cross-modal perception that requires the integration of information from distributed neuronal assemblies. Taken together, communication via alpha coherence may be the most plausible neuromarker of inter-group variability at the network level bearing the signature of integration.

### Multisensory node operational at intermediate time-scale renders communication effective between sensory specific nodes operating at slow and fast time scales

Neural mass models that operate to explain mesoscopic and macroscopic features of dynamics in EEG/ MEG data offers an attractive tool to understand the underlying neural mechanisms (Silva et al., 1974; Jansen & Rit, 1995; David & Friston, 2003). The time scale of processing in our computational model is most disparate for the auditory and visual node, with auditory the fastest and visual the slowest (Rosen & Howell, 2011; Williams et al., 2004). Without the presence of an intermediate time-scale, only alpha coherence is sustained by the neural mass model within biologically relevant parameter regimes (**Figure 6**). On introduction of another neural mass operating at intermediate time-scale, synchronization in the gamma band emerges because higher dimensionality of the resultant dynamical system allows creation of another mode of communication exchange. Hence, our model suggests that emergence of gamma band coherence could be attributed to the communication between areas of auditory and visual cortices gated through multisensory areas such as STS (**Figure 7**). Our suggestion is in line with earlier observations of visual stimuli modulating auditory perception either directly resulting in alpha band coherence (Kayser et al., 2008) or indirectly via higher order regions resulting primarily in gamma band coherence (Maier et al., 2008; Kayser & Logothetis, 2009).

Drawing from the model, inter-individual differences primarily emerge from the differences in the A-V coupling. A stronger A-V coupling implies a lesser mediation via the multisensory node (stronger A-M and V-M) leading to weaker integration which is akin to rare perceivers. Conversely, lesser A-V coupling implies a larger involvement of the multisensory node (stronger A-M and V-M) in mediating the integration of the sensory inputs that is characteristic of frequent perceivers. Findings from our model corroborates with the earlier findings that pinpoint the significance of functional connectedness of the multisensory regions like STS to sensory specific, frontal and parietal regions in facilitating cross-modal perception (Keil et al., 2012).

There are a number of qualifications that attend this work and its interpretation. In brief, our exploration of different connectivity architectures was limited to extrinsic (i.e., between node) coupling differences. These differences were both in terms of the arrangement of the connections – and connection strengths. However, we did not consider intrinsic connectivity; namely, within node connections. The role of intrinsic connectivity in accounting for inter-individual differences is something that we hope to explore in the future. This is a potentially important avenue of enquiry, given the sensitivity of global dynamics to subtle changes in intrinsic connectivity and excitability (Freyer et al., 2012; Roy et al., 2014; Gilson et al., 2016). This is particularly prescient from the point of view of predictive coding models of multisensory integration. In this setting, the precision assigned to various sources of information (i.e., prediction errors) is thought to be mediated by intrinsic excitability (or self-inhibition) (Bhatt et al., 2016; Brown & Friston, 2012; Pinotsis et al., 2014). We also note, in contrast to the neural mass models used in dynamic causal modeling (Bastos et al., 2012; Bastos et al., 2015; Litvak et al., 2015), that the designation of forward and backward is based purely upon the distinction between excitatory and inhibitory connectivity. In biologically plausible models this designation usually rests upon laminar specificity (Markov et al., 2013; Shaw et al., 2019). However, our neural mass model did not distinguish between different excitatory and inhibitory populations in different cortical layers. It would be interesting to reproduce the analysis reported above using nuanced models of the sort used in in dynamic causal models based on the canonical microcircuit. Provisional work in this area has already considered A/V sensory integration (Li et al., 2019); however, it has not yet addressed the cardinal issue of inter-individual variability.

### Dynamic coupling between audio and visual cortices underlie perceptual variability

Most importantly, we validated our predictions from the dynamical model to the empirical estimates of source-level functional connectivity. Our pairwise connectivity between the parcels (**Figure 8**) show increased connectivity between the auditory and visual parcels in rare than frequent perceivers. Interestingly, we observed this increase only in the left hemisphere which corroborates the earlier findings accentuating left laterality in multisensory speech perception (Beauchamp, 2010; Nath & Beauchamp, 2012; Keil et al., 2012). Notably, for our source reconstruction, we used a form of beamforming (DICS) to identify sources that were likely to contribute the most power in sensor space. While this was sufficient for our purposes, this kind of source reconstruction has to be used carefully when subsequently measuring coherence among sources. This is because the scheme used to identify or estimates sources suppresses coherent activity between sources. It will be interesting to perform a more detailed source reconstruction that embodies our prior knowledge about the key nodes or sources involved in the McGurk effect; for example, using equivalent current dipole reconstructions that are guided by the functional magnetic resonance imaging (fMRI) literature in this area.

The analysis pipeline (**Figure 1**), can be generalized easily to other studies of complex perceptual phenomena. For example, alpha and/or gamma coherences have been observed in AV perception studies involving speech phrases (Doesburg et al., 2008), natural scenes (Kayser et al., 2008) and also artificially generated A-V looming signals (Maier et al., 2008). Modulations in A-V interactions that explains differences in McGurk susceptibility in our study could also explain the increase in alpha phase consistency observed during natural AV scenes in rhesus monkeys (Kayser et al., 2008). Our model demonstrates that increase in the interaction between neural nodes operating at fast and slow time scales via a node operating at an intermediate time scale results in increased gamma and decreased alpha and beta band coherence. However, a slight tweak in the model topology elicits coherence modulations in varied frequency bands. A basic understanding that multifrequency coherent states of the network emerges from their difference in time scales can be used to explain a large repertoire of empirical observations.

Although our model of three interacting neural masses with different time-constants that generate band specific coherences can come across as a canonical model of neuronal network dynamics, although, certain limitations exist. Multi-parametric and unbounded nature of the parameter space results in myriads of dynamics including chaos (Stefanescu & Jirsa, 2008). Therefore, such models cannot be used to directly fit the data using optimization techniques. and should be limited to use as a phenomenological or minimalistic model in providing mechanistic insights into many findings (Fries, 2015; Engel et al., 2012),-both in normal and clinical population. Also, this kind of neural mass model can be contrasted with the neural mass models used in dynamic causal modelling, which have the fixed point attractors and can be used to fit observed EEG time-series or spectra.

## Abbreviations

EEG: Electroencephalogram
AV: Audio-visual
STS: Superior temporal sulcus
LGN: Lateral geniculate nucleus
MGN: Medial geniculate nucleus
DICS: Dynamic imaging of coherent sources
BEM: Boundary Element Method
AAL: Automated Anatomical Labelling
HR: Hindmarsh-Rose
SOC: Superior Occipital Cortex
MOC: Medial Occipital Cortex
IOC: Inferior Occipital Cortex
STG: Superior temporal gyrus
MEG: Magnetoencephalogram
fMRI: functional magnetic resonance imaging

## Acknowledgements

This study was supported by NBRC Core funds, Ramalingaswami Fellowships (Department of Biotechnology, Government of India) to DR (BT/RLF/Re-entry/07/2014) and AB (BT/RLF/Re-entry/31/2011) and Innovative Young Biotechnologist Award (IYBA) to AB (BT/07/IYBA/2013). DR was supported by SR/CSRI/21/2016 extramural grant from the Department of Science and Technology (DST) Ministry of Science and Technology, Government of India.

